# DNA-binding specificity recognition from predicted homologous protein-DNA structures

**DOI:** 10.64898/2026.06.16.732781

**Authors:** Wenwu Zeng, Haitao Zou, Xiaoyu Li, Yaoyu Liu, Liwen Xu, Xiaoqi Wang, Shaoliang Peng

## Abstract

Predicting protein DNA-binding specificity is essential for understanding gene regulation and disease mechanisms. Existing deep learning methods typically infer specificity from a single protein-DNA complex structure, which limits their ability to capture the diverse geometric patterns underlying protein-DNA recognition. Homologous protein-DNA interfaces provide complementary structural evidence and richer geometric features related to interatomic interactions. To address the limited diversity and coverage of experimentally determined complexes, we constructed a large-scale library of predicted homologous protein-DNA complex structures. Building on this resource, we propose HomoDSP, a template-retrieval-based framework for accurate DNA-binding specificity prediction. Benchmark evaluations and validation on newly released JASPAR 2026 samples indicate that HomoDSP outperforms existing methods in both accuracy and generalization, with particularly substantial gains on high-error samples. Moreover, this performance is largely retained when AlphaFold3-predicted complex structures are used as input. Template- and residue-level interpretability analyses suggest that HomoDSP improves prediction by focusing on DNA-affinity residues across multiple homologous templates. Finally, universal Protein Binding Microarrays evaluations on AI-designed DNA-binding proteins show that HomoDSP rescues a baseline failure mode in which the baseline method produces incorrect predictions because of training-set bias. Together, these results support the use of homologous template interfaces as informative structural priors for decoding protein DNA-binding specificity.

## Introduction

Sequence-specific DNA-binding proteins (DBPs) play central roles in gene regulation by recognizing defined DNA sequence patterns in regulatory elements. Among them, transcription factors (TFs) establish cell-type-specific transcriptional programs by selectively occupying genomic loci and recruiting regulatory cofactors [1–4]. Perturbations of DNA recognition can rewire gene expression and contribute to disease. Accurately characterizing DNA-binding specificity (DBS) is essential for understanding gene regulation, interpreting regulatory variants, and engineering synthetic DBPs [5–7].

DBS is commonly represented by position weight matrices (PWMs), which summarize the base preferences of a DBP across aligned binding sites. These profiles are widely used for motif discovery, TF binding-site prediction, regulatory variant annotation, and synthetic biology applications. Large-scale experimental assays such as HT-SELEX, DAP-seq, and ChIP-seq have generated valuable specificity profiles for many TFs. However, these experimental measurements remain costly, condition-dependent, and unevenly distributed across species and protein families. *In vitro* assays such as universal protein-binding microarrays (uPBMs) more directly measure intrinsic DNA-binding preferences but cover only a fraction of natural DBPs. Consequently, many orphan DBPs, non-model organism TFs, and newly identified homologs still lack reliable specificity profiles. This motivates computational approaches that can infer DBS directly from protein sequence and structure [8]. Furthermore, accurate and generalizable DBS prediction with PWM outputs provides an important high-throughput screening tool for the rapidly developing field of AI-driven DBP design [9–13].

Protein-DNA complex structures provide a direct view of the molecular interactions underlying DNA recognition. They show how amino acid side chains contact bases, phosphate groups, and the DNA backbone, and how protein shape complements local DNA geometry. Recent geometric deep learning methods such as DeepPBS [14] and NA-MPNN [15] have shown that such structural information can be used to predict DBS. Despite these advances, current structure-based specificity predictors are constrained by the limited and redundant nature of available protein-DNA complex structures. Experimentally determined complexes in the PDB cover only a small subset of DBP families, and many structures are highly similar. Moreover, a single protein-DNA complex captures only one bound DNA sequence and one recognition geometry, whereas DBS describes a broader landscape of tolerated and disfavored DNA sequences. This creates a fundamental gap: the structural state observed in one complex is informative but may not be sufficient to represent the family-level conservation and divergence of DNA recognition. In particular, homologous DNA-binding domains (DBDs) often preserve a global conformation while differing in local DNA-contacting residues, scaffold context, and interface geometry. These variations may contain specificity-preserving or specificity-shifting signals that are difficult to extract from a single query structure alone. **We hypothesize that homologous protein-DNA complexes provide a complementary source of recognition evidence for DBS prediction**. Rather than treating each protein-DNA structure as an isolated example, we view related complexes as a homologous recognition landscape. Within this landscape, conserved structural scaffolds define the overall DNA-binding mode, whereas local variations in contact residues, side-chain environments, and DNA-interface geometry may provide information about how specificity is maintained or altered across homologous proteins. This perspective is distinct from simply transferring the motif of the nearest homolog. Instead, it asks whether a set of homologous protein-DNA interfaces can provide structure-aware evidence that improves specificity inference beyond a single query complex.

The primary requirement is a sufficiently large database of homologous complexes that extends beyond the limited set of experimentally solved structures. The emergence of accurate biomolecular structure prediction methods, particularly AlphaFold3 [16] (AF3), makes this possible. A key challenge is how to construct biologically meaningful protein-DNA pairs at scale. **DBPs with similar DBDs tend to exhibit analogous DNA-binding preferences (Fig. 2a) [17–19]**. We reasoned that experimentally determined protein-DNA complexes provide reliable anchors for DNA recognition. Starting from known protein-DNA complexes, we identify DBDs and search sequence databases for homologous proteins sharing similar domains. These homologous proteins are then paired with the DNA molecules from the anchor complexes and modeled with Protenix [20] (a re-implementation of AF3) to generate predicted homologous protein-DNA complexes as the template database (Fig. 1a). **Mutations in individual residues can lead to conformational changes at the protein-DNA binding interface [21–23]**. **Such geometric variation provides interaction-relevant features that may be useful for DBS prediction**. We do not assume that every homolog necessarily binds the paired DNA *in vivo*. Instead, these predicted complexes serve as structurally plausible recognition contexts that preserve the global DNA-binding mode while introducing natural variation in protein sequence, scaffold context, and interface geometry.

**Figure 1.**
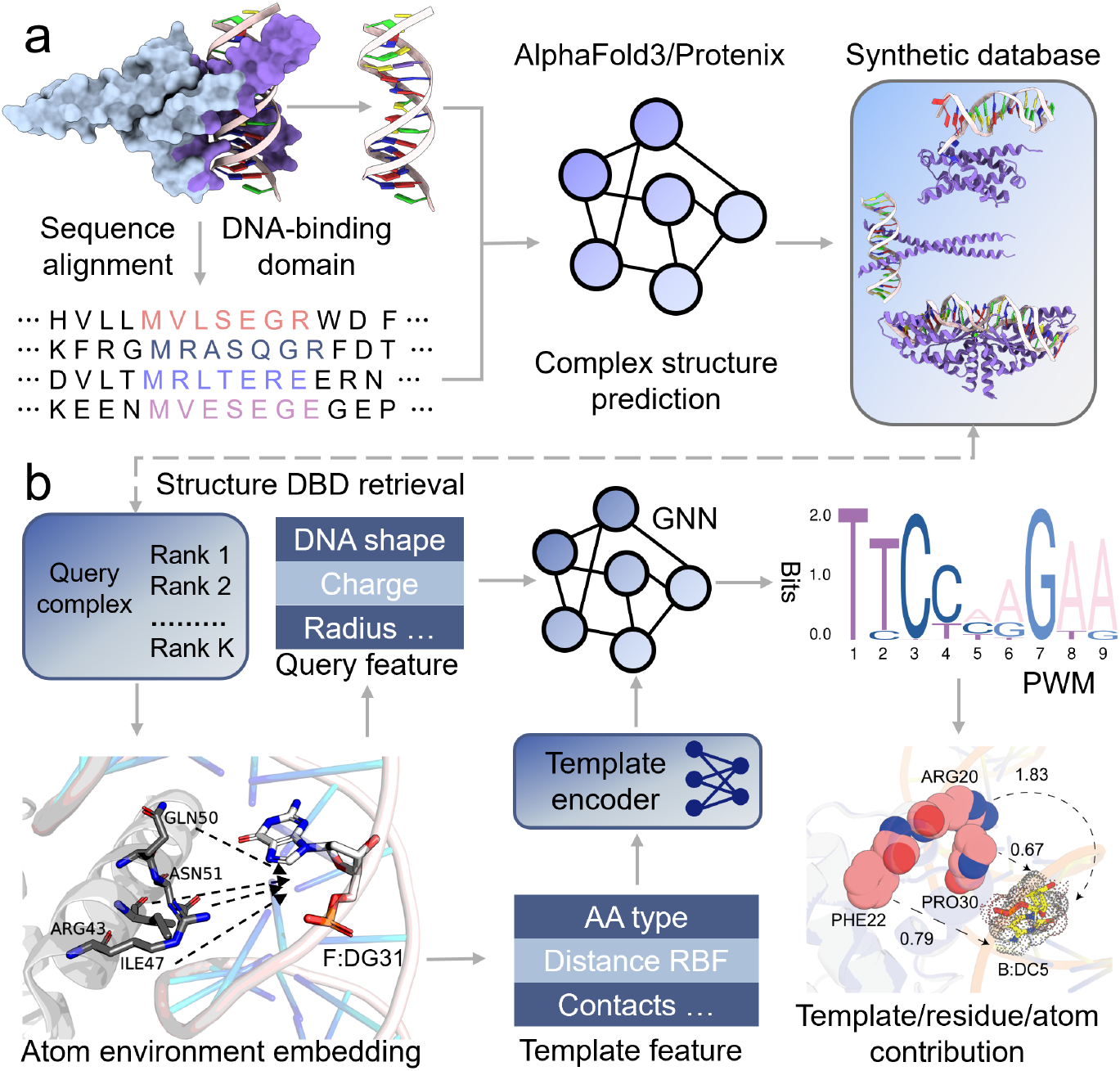
Homology-derived structure templates for DBS prediction. **a**, DBP database synthesis. DBD sequences are extracted from native protein-DNA complexes and used to retrieve homologous protein domains from sequence databases. Each homologous sequence is paired with the target DNA sequence from the source complex and submitted to a complex structure predictor. **b**, DBS prediction. For each query complex, Foldseek is used to retrieve homologous complex templates from the synthetic structure database. Atomic environments of the query and template complexes are encoded as geometric features and integrated by a geometric neural network to predict the DNA-binding PWM. Finally, gradient-based attribution estimates template/residue/atom-level contributions.

**Figure 2.**
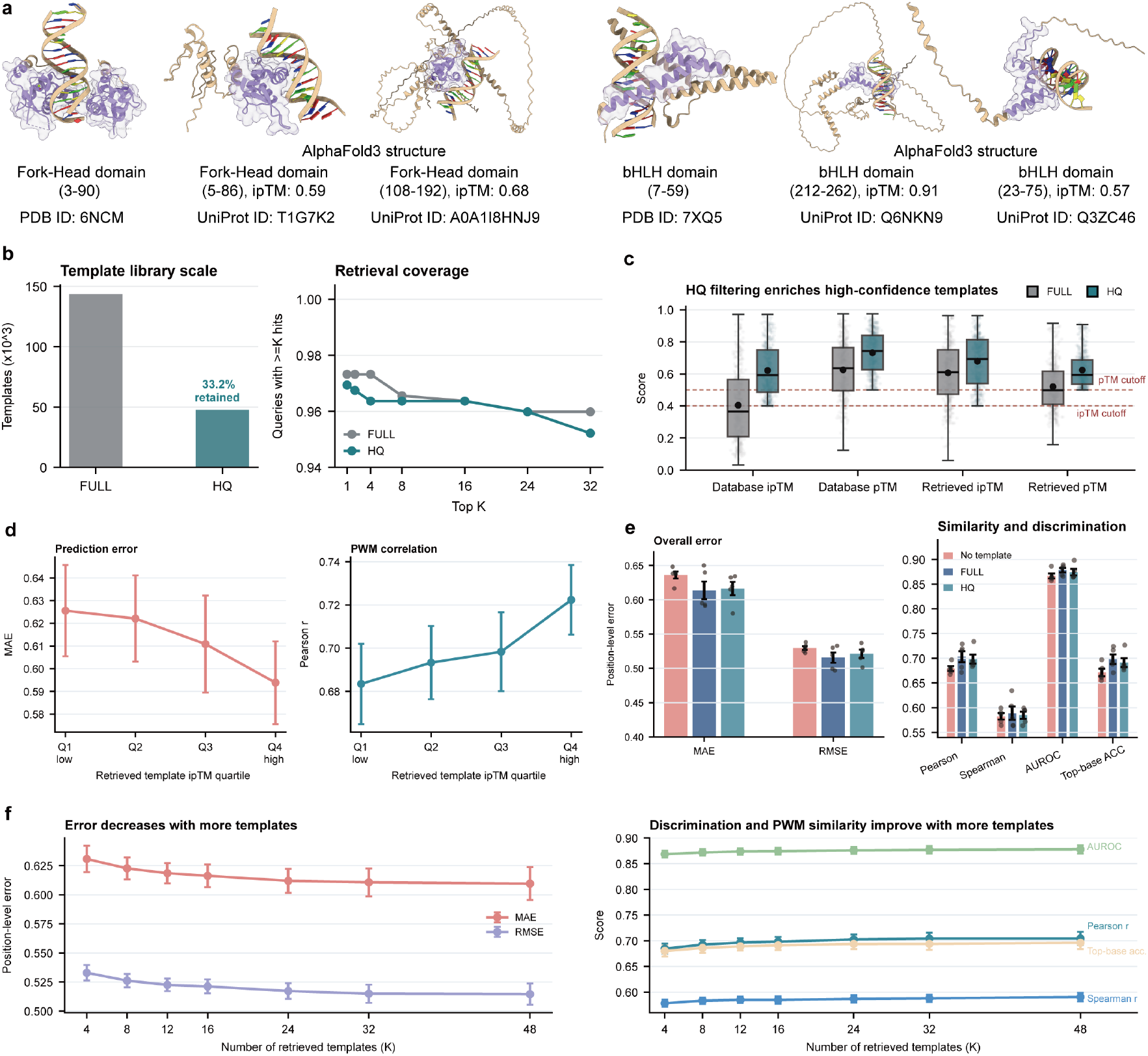
Validity of homologous protein-DNA complex templates for DBS prediction. **a**, Examples of homologous complex templates. Experimentally determined protein-DNA complexes from the PDB are compared with AlphaFold3-predicted complexes formed by pairing homologous protein sequences with the same DNA molecule. Purple highlights DBDs, which retain plausible binding geometries across different sequence contexts. **b**, Database scale and retrieval coverage. The HomoDB-FULL database contains approximately 1.45 × 10^5^ predicted complexes, and the HomoDB-HQ subset retains 33.2% after confidence filtering while preserving high top-*K* retrieval coverage. **c**, HomoDB-HQ filtering enriches high-confidence templates, as shown by ipTM and pTM distributions for all and Foldseek-retrieved templates. Dashed lines mark filtering thresholds. **d**, Higher retrieved-template ipTM is associated with lower PWM prediction error and higher PWM correlation. **e**, Cross-validation performance comparison of models without templates, with templates from HomoDB-FULL, and with HomoDB-HQ templates. **f**, Increasing the number of retrieved templates improves prediction performance, with gains saturating at larger *K*. Error bars denote the standard error of the mean across five folds.

We introduce HomoDSP, a homology- and structure-guided framework for predicting DBS from homologous protein-DNA structures. Given a query protein-DNA complex, HomoDSP retrieves homologous structural templates from the template database and integrates their interface features with the query structure. The model consists of geometric neural networks that process both the query complex and the retrieved homologous templates, enabling DBS prediction with PWM outputs from structure-aware representations of protein-DNA recognition (Fig. 1b). By design, HomoDSP aims to exploit not only the local geometry of the query structure but also the conserved and divergent interface patterns present across homologous complexes. This framework addresses a central limitation of single-structure specificity prediction: one bound complex provides only a partial view of the recognition landscape. By incorporating homologous templates, HomoDSP uses additional recognition-compatible protein-DNA interfaces to inform base-preference prediction. More broadly, HomoDSP tests whether conserved and variable interface patterns across homologous DBPs contain informative signals for DBS prediction.

We systematically assessed HomoDSP across multiple settings. Five-fold cross-validation (CV) on benchmark datasets showed that homologous structural templates consistently improve DBS prediction and outperform existing structure-based DBS predictors. Evaluation on 33 newly released JASPAR 2026 samples further supported the generalizability of HomoDSP. Family-level analyses indicated broad performance gains across most major DBP families, while sample-level comparisons showed that these gains are concentrated in high-error cases where single-structure predictors exhibit high baseline error. Gradient-based attribution further indicated that HomoDSP improves individual motif-column predictions by leveraging canonical DNA-contacting and interface-stabilizing residues from homologous template complexes, with local geometry providing the dominant predictive signals. On AF3-predicted complexes, HomoDSP incurred only a modest performance loss for high-confidence structures and still outperformed the baseline, supporting its applicability in a sequence-to-structure-to-specificity prediction setting. Finally, uPBM evaluation on AI-designed DBPs showed that HomoDSP can correct a baseline failure mode associated with training-set-dependent local-interface ambiguity by using heterogeneous homologous template geometries to distinguish locally similar protein-DNA environments with different nucleotide preferences. Together, these findings suggest that conserved and variable interface patterns across homologous complexes contain informative signals for DBS prediction. By enabling accurate, interpretable, and scalable DBS prediction, HomoDSP provides a practical framework with the potential to accelerate systematic studies of protein-DNA interactions.

## Results

### The HomoDSP framework

The HomoDSP framework (Fig. 1) consists mainly of two parts: **(1) construction of the predicted homologous complex database** (see Section “The predicted homologous complex database”). Given a native DBP, homologous protein sequences are first retrieved and paired with the same DNA sequences to generate predicted protein-DNA complexes. By traversing the PDB database, we obtained a large-scale, searchable homologous DBP template database named HomoDB. **(2) template-enhanced geometric neural networks for DBS prediction** (see Section “Template-enhanced geometric neural networks” and Appendix A). For each query complex, homologous templates are retrieved, aligned, and converted into atom-level interface features. HomoDSP then integrates the query protein-DNA geometry with template-derived interface information through a geometric neural network to predict the corresponding PWM. The evaluation metrics are described in Appendix B, and the datasets used in this study are described in Section “Evaluation datasets”.

### Validity of homologous template databases

We first evaluated whether predicted homologous protein-DNA complexes provide useful structural evidence for DBS prediction. Using HomoDB-FULL as the template library, HomoDSP consistently outperformed the template-free variant in 5-fold CV on the benchmark dataset (Fig. 2e), indicating that homologous templates provide complementary information beyond the query complex alone. Prediction performance was also associated with template confidence: queries retrieving templates with higher ipTM scores showed lower PWM prediction error and higher PWM correlation (Fig. 2d). This suggests that more reliable predicted protein-DNA interfaces generally provide more informative template features for DBS prediction.

We next assessed the high-quality subset, HomoDB-HQ. Confidence filtering retained 33.2% of the full database while only modestly reducing top-*K* retrieval coverage (Fig. 2b). HomoDB-HQ achieved performance comparable to HomoDB-FULL (Fig. 2e). This result suggests that template confidence is an important but not exclusive determinant of prediction accuracy. On this benchmark, the templates retrieved from HomoDB-FULL were already of sufficient quality (Fig. 2c), and the greater diversity of the full database may provide additional geometric information that compensates for lower average confidence.

Finally, we examined the effect of the number of retrieved templates. Increasing *K* improved prediction performance, particularly when moving from small template sets to moderate template coverage, but the gains gradually saturated at approximately *K* = 32 (Fig. 2f). Together, these analyses support the validity of the predicted homologous template library and show that HomoDSP benefits from both confident protein-DNA interface predictions and diverse homologous structural contexts.

### Comparison with existing structure-based predictors

We compared HomoDSP (top-16 templates) with two state-of-the-art structure-based DBS predictors, DeepPBS and NA-MPNN (re-implementation using the same dataset), on the 5-fold CV benchmark and 33 newly released JASPAR2026 samples. HomoDSP achieved the best overall performance across most error and similarity metrics, reducing PWM prediction error while improving motif similarity and top-base accuracy relative to both baselines (Fig. 3a,b). These results indicate that homologous structural templates provide complementary recognition information beyond the single-complex representations used by existing methods.

**Figure 3.**
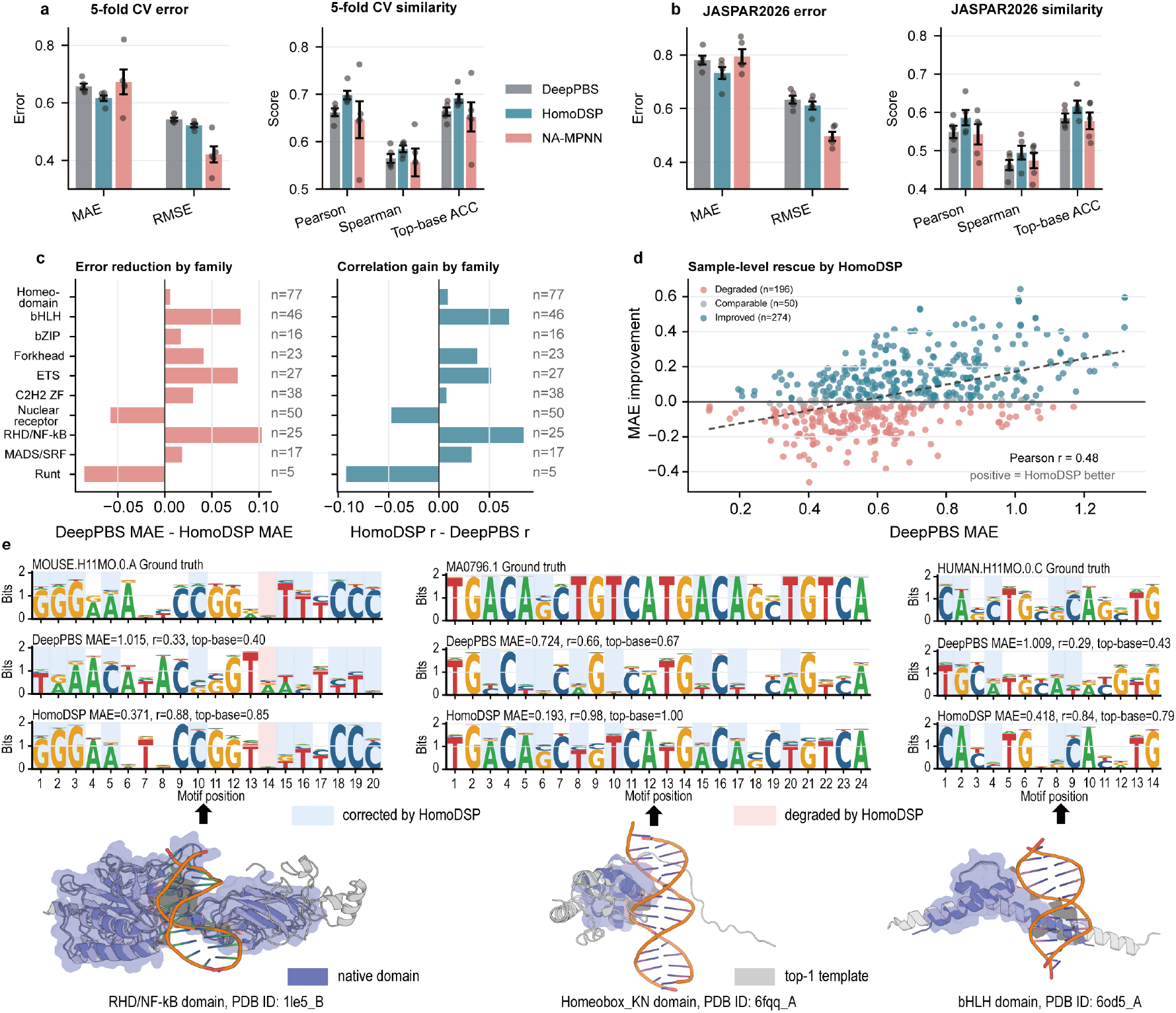
HomoDSP improves structure-based prediction of DBS over existing methods. **(a-b)**, 5-fold CV and JASPAR2026 evaluation results showing that HomoDSP reduces PWM prediction error and increases similarity to ground-truth motifs compared with DeepPBS and NA-MPNN (re-implementation on the same dataset). Bars show mean performance across 5 folds; points denote individual folds. **c**, Family-level gains in MAE and PCC. Positive values indicate improved performance by HomoDSP, demonstrating consistent benefits across most TF families. **d**, Sample-level improvements over DeepPBS. HomoDSP preferentially rescues high-error cases, as indicated by dense points in the upper right. **e**, Case studies showing template-guided rescue of motif positions. HomoDSP better recovers ground-truth sequence logos by using homologous protein-DNA complex templates that provide local interface geometry.

We observed that HomoDSP achieved a lower MAE but a slightly higher RMSE than NA-MPNN in the cross-validation benchmark. This pattern suggests that HomoDSP reduces the typical prediction error but has a heavier error tail. In other words, homologous templates improve predictions for most samples, whereas a small subset of high-error cases showed larger deviations. By contrast, NA-MPNN produced fewer extreme errors but weaker average improvement. Family-level analysis showed that HomoDSP produced consistent gains in MAE and Pearson correlation across most major transcription factor families (Fig. 3c). The main exception was the nuclear receptor family (*n* = 50), for which HomoDSP showed degraded performance. This degradation is consistent with lower template quality in this family, as suggested by the family-level template quality analysis (Fig. A1). These results indicate that the benefit of homologous templates is broadly observed but remains family-dependent.

We next examined whether HomoDSP preferentially improves high-error samples. **Sample-level comparisons against DeepPBS showed a positive correlation between DeepPBS baseline error and HomoDSP improvement (Pearson** *r* = 0.48**; Fig. 3d)**, **indicating that HomoDSP gains are concentrated in cases where single-structure prediction is less reliable**. Representative examples further illustrate this behavior (Fig. 3e). In `1le5_B` (RHD/NF-*κ*B domain), HomoDSP substantially improved the recovery of the C/G-rich repetitive pattern that was poorly captured by DeepPBS. Similar template-guided corrections were observed for `6fqq_A` (Homeobox_KN domain) and `6od5_A` (bHLH domain), although the latter also shows that template information can occasionally degrade individual positions. Overall, these results indicate that HomoDSP improves structure-based DBS prediction not merely by reducing average error, but also by using homologous protein-DNA templates to recover motif positions that are ambiguous from the query complex alone.

However, we also observed mild degradation for a subset of easy samples with low DeepPBS error. In these cases, the query complex may already contain sufficient information to determine the motif, and additional homologous templates can introduce unnecessary variance or dilute a clear local recognition signal, particularly when the retrieved templates represent related but slightly divergent specificity states. This suggests that template information is most beneficial when the query structure alone is ambiguous, but may be less helpful when the intrinsic structural signal is already strong.

### Template attribution explains HomoDSP rescue of baseline failures

We performed gradient-based interpretability analysis to understand how HomoDSP improves DBS prediction on baseline failures (Fig. 4). We focused on three representative cases in which DeepPBS failed to recover the correct base preference, whereas HomoDSP corrected the corresponding motif column.

**Figure 4.**
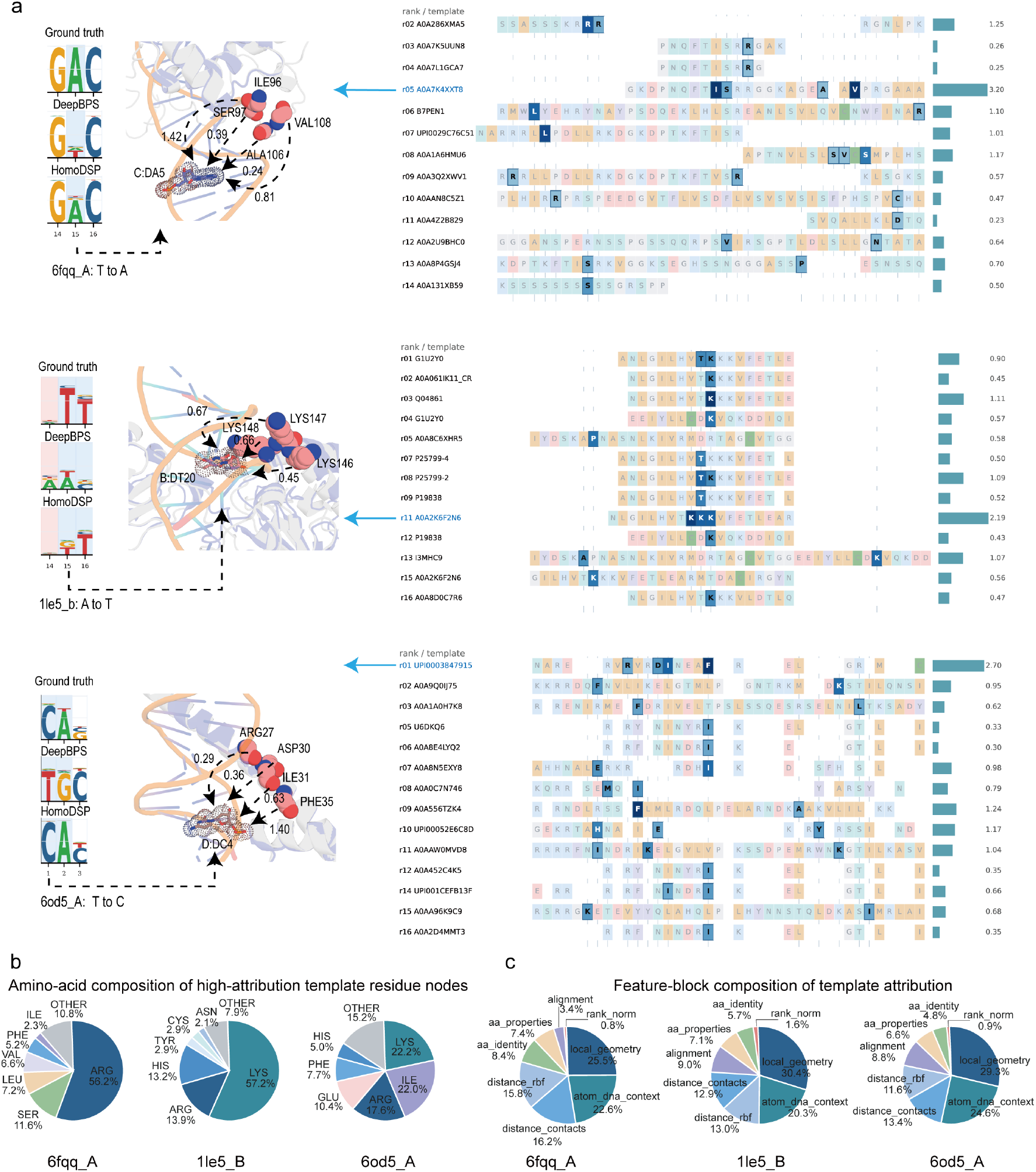
Template/residue-level attribution analysis explains HomoDSP improvements. **a**, Template/residue-level attribution for three representative examples where HomoDSP improves DBS prediction over DeepPBS. High-attribution residues near the target base are shown as spheres; numbers denote residue contributions. The right panel shows residue-level contributions across retrieved templates. High-attribution residues are highlighted in blue ↑. **b**, Amino-acid composition of high-attribution template residues. Basic residues Arg and Lys, together with hydrophobic residue Ile, contribute most strongly, consistent with their roles in DNA contact and interface stabilization. **c**, Feature-block composition of template attribution. Local geometry and atom-level DNA-context features dominate, indicating that HomoDSP mainly uses structure-aware protein-DNA interface features rather than sequence similarity alone.

Across the three representative cases, HomoDSP corrected nucleotide preferences that were mispredicted by DeepPBS by leveraging high-attribution residues from homologous template interfaces (Fig. 4a). In `6fqq_A`, HomoDSP rescued the target position from an erroneous T prediction to the ground-truth A preference, with recurrent Arg-containing positions contributing across multiple retrieved templates. In `1le5_B`, the correction from A toward the ground-truth T preference was mainly associated with a Lys-rich interface cluster near the target base, including `LYS146`, `LYS147` and `LYS148`. In `6od5_A`, HomoDSP corrected a T-to-C error, with attribution distributed across residues such as `ARG27`, `ASP30`, `ILE31` and `PHE35`. These examples indicate that HomoDSP does not rely on a single nearest template, but instead integrates residue-level evidence from multiple homologous interfaces to disambiguate local protein-DNA recognition patterns.

We next summarized the amino-acid composition of high-attribution template residues (Fig. 4b). Basic residues, especially Arg and Lys, accounted for the largest contributions in `6fqq_A` and `1le5_B`, consistent with their well-established roles in contacting the negatively charged DNA backbone and stabilizing protein-DNA interfaces. In `6od5_A`, hydrophobic residues such as Ile also contributed strongly, suggesting that local packing and recognition-helix stabilization can provide additional specificity-relevant information beyond direct electrostatic DNA contacts.

Finally, feature-block attribution showed that local geometry and atom-level DNA-context features dominated the template contribution across the three examples (Fig. 4c). In contrast, sequence identity and alignment-related features contributed less. These results indicate that HomoDSP improves DBS prediction primarily by extracting structure-aware interface information from homologous protein-DNA templates. Together, the interpretability analysis supports the view that homologous templates help disambiguate local protein-DNA environments that appear similar in a single query structure but are associated with different nucleotide preferences.

### Performance on AF3-predicted structures

In the post-AF3 era, predicted protein-DNA complex structures provide an important structural basis for large-scale DBS prediction. We therefore evaluated HomoDSP on Protenix-predicted protein-DNA complex structures for the 5-fold CV benchmark (Fig. 5). This analysis was designed to assess whether HomoDSP remains effective when experimental structures are replaced with predicted complexes.

**Figure 5.**
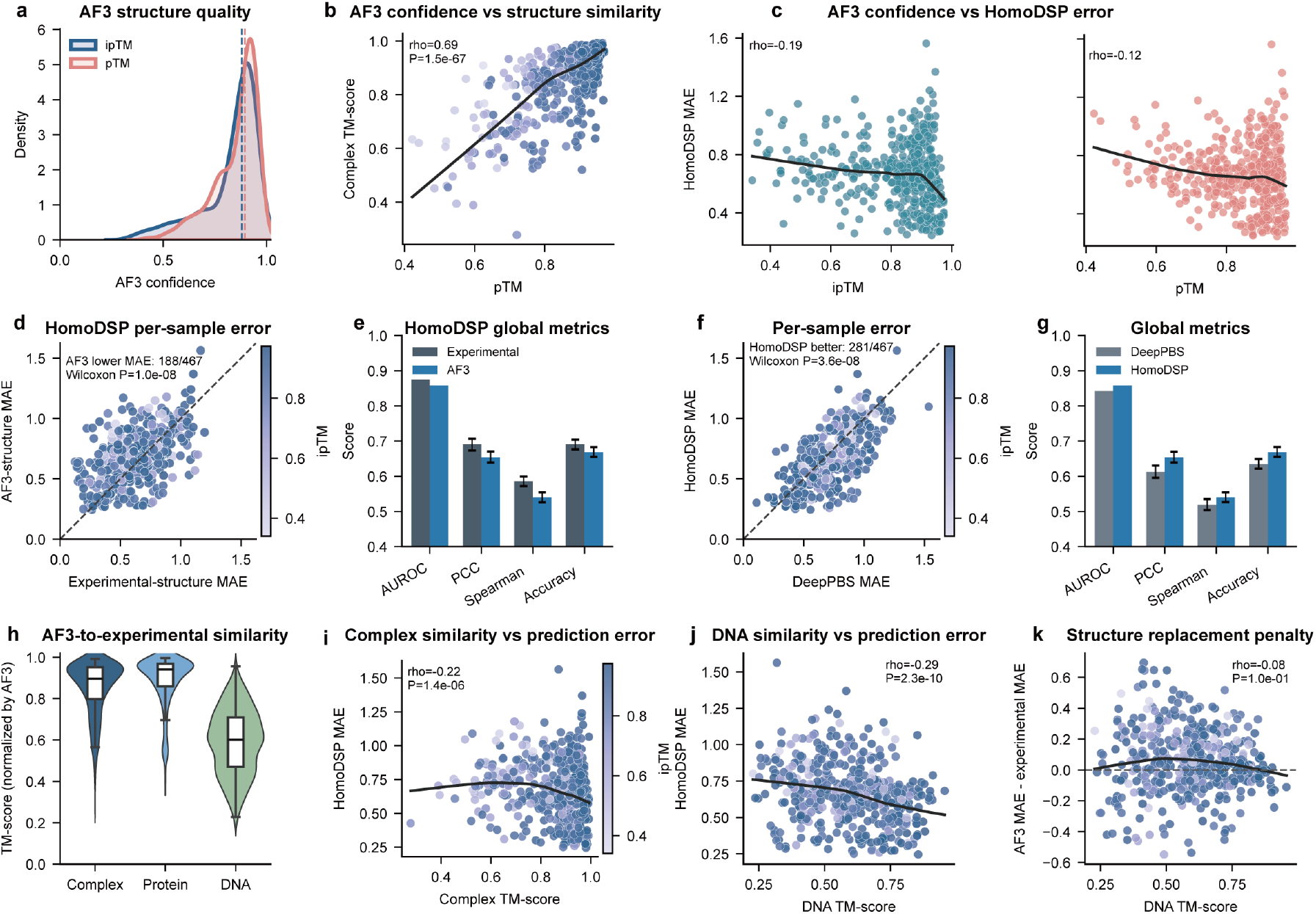
Evaluation on predicted protein-DNA complex structures. **(a)** Distribution of AF3 confidence scores, i.e., ipTM and pTM. **(b)** Relationship between pTM and complex TM-score, showing whether AF3 global confidence reflects similarity to the experimental complex structure. **(c)** Relationship between AF3 confidence and HomoDSP prediction error. Higher ipTM or pTM generally corresponds to lower PWM MAE. **(d-e)** Comparison between HomoDSP predictions using experimental structures and AF3-predicted structures. Per-sample MAE and global metrics quantify the performance change caused by replacing experimental structures with AF3 models. **(f-g)** Comparison between DeepPBS and HomoDSP when both use the consistent AF3-predicted structures. HomoDSP achieves lower per-sample error and better global metrics. **(h)** TM-score distributions for whole-complex, protein-only, and DNA-only alignments between AF3 and experimental structures. **(i-j)** Correlation between structural similarity and prediction reliability. Higher complex or DNA TM-score is associated with lower HomoDSP MAE. **(k)** Scatter plot showing the relationship between DNA TM-score and the structure replacement penalty, defined as the difference in HomoDSP MAE between AF3-predicted and experimental structures.

**First, the predicted complexes showed generally high confidence**. Both ipTM and pTM were concentrated at high values (Fig. 5a), consistent with the strong structural conservation of many DBDs. Moreover, pTM was positively correlated with the complex-level TM-score (calculated by US-align [24]) between predicted and experimental structures (Spearman *ρ* = 0.69, *P* = 1.5 × 10^−67^; Fig. 5b). **Second, AF3 confidence was informative for downstream DBS prediction reliability**. HomoDSP prediction error showed weak but consistent negative correlations with both ipTM and pTM (Spearman *ρ* = −0.19 and −0.12, respectively; Fig. 5c). Although these correlations were modest, they indicate that higher-confidence predicted interfaces tend to yield lower PWM prediction error. Thus, AF3 confidence can provide a practical quality indicator for large-scale DBS prediction, but it is not sufficient to fully determine downstream accuracy. **Third, replacing experimental structures with AF3-predicted structures caused only a modest performance loss**. Per-sample comparison showed that HomoDSP predictions from AF3 structures were close to those from experimental structures, although AF3-based inputs produced slightly higher MAE for a subset of samples (Fig. 5d). Consistently, global metrics showed only mild degradation in AUROC, PCC, Spearman correlation, and top-base ACC (Fig. 5e). These results suggest that HomoDSP is robust to realistic structural perturbations introduced by predicted complexes. **Fourth, HomoDSP still outperformed the baseline method when AF3-predicted structures were used**. When both methods used consistent AF3-predicted structures as input, HomoDSP achieved lower per-sample error for most samples and improved global similarity metrics relative to DeepPBS (Fig. 5f,g). This result indicates that the benefit of homologous template augmentation is largely preserved even when the query structure is computationally predicted rather than experimentally determined. **Fifth, structural similarity analysis showed that protein and complex structures were generally well recovered, whereas DNA geometry was more variable**. AF3-predicted complexes showed high similarity to experimental structures at the protein and whole-complex levels, but DNA-only TM-scores were substantially lower and more broadly distributed (Fig. 5h). Higher complex and DNA TM-scores were associated with lower HomoDSP MAE (Fig. 5i,j), suggesting that both global complex accuracy and DNA geometry contribute to prediction reliability. However, the AF3-induced replacement penalty showed only a weak relationship with DNA TM-score (Fig. 5k), indicating that DNA structural deviation alone cannot fully explain the observed performance changes.

Together, these results indicate that HomoDSP can effectively operate on AF3-like predicted protein-DNA complex structures. Although prediction reliability remains influenced by structural confidence and local DNA/interface geometry, the modest performance loss and sustained improvement over DeepPBS support the use of predicted complexes for large-scale DBS prediction.

### uPBM validation on AI-designed DBPs

We next evaluated whether HomoDSP improves DBS prediction for AI-designed DBPs using experimentally measured uPBM data from Glasscock *et al*. [13]. For each designed DBP, we compared the predicted PWM with uPBM-derived 7-mer E-scores by scanning all possible 7-bp PWM windows and selecting the window with the highest Spearman correlation with the experimental measurements.

Across the seven AI-designed DBPs, HomoDSP showed its largest gain on DBP48 (Fig. 6a). Although performance varied across designs, the Spearman correlation for DBP48 increased from 0.14 for DeepPBS to 0.35 for HomoDSP. At the 7-mer level, HomoDSP also assigned higher percentiles to a larger fraction of experimentally enriched uPBM 7-mers than DeepPBS (Fig. 6b), indicating better recovery of the experimental ranking of DBP48 binding preferences.

**Figure 6.**
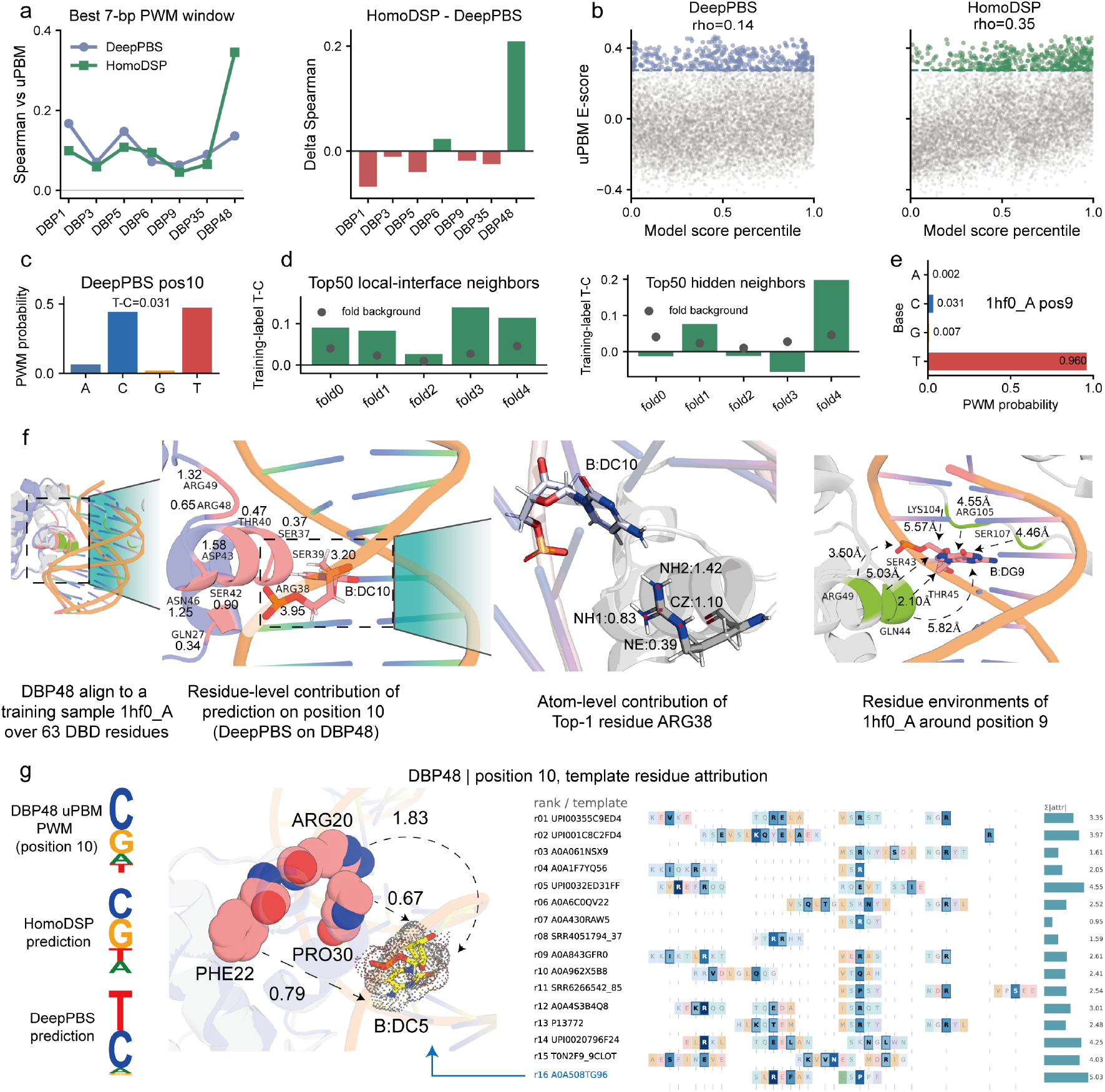
HomoDSP resolves a baseline failure mode in AI-designed DBPs. **a**, Left, Spearman correlation for DeepPBS and HomoDSP. Right, per-protein correlation gain of HomoDSP over DeepPBS. **b**, DBP48 score calibration against uPBM measurements. All 7-mers are ranked by model-score percentile and colored points denote top model predictions. **c**, DeepPBS prediction at DBP48 position 10. **d**, Training-neighborhood analysis for DBP48 position 10. Top-50 training positions were retrieved by either a local protein-DNA interface descriptor (or the DeepPBS hidden representation) before PWM readout. Bars show mean training-label *P* (*T*) − *P* (*C*) for each checkpoint; gray points show fold-specific background. **e**, PWM label of representative hidden neighbor `1hf0_A` position 9, a neighbor of DBP48 position 10, showing strong T preference. **f**, Structural and attribution analysis of the DeepPBS error on DBP48 position 10. DeepPBS integrated-gradient attribution for the T-over-C prediction highlights residues near the queried base. Atom-level attribution concentrates on the ARG38 guanidinium group, indicating that the erroneous prediction is driven by positively charged DNA-contact atoms. The corresponding `1hf0_A` environment contains similar basic/polar residues, suggesting that DeepPBS maps DBP48 position 10 to a structurally related but T-favoring training neighborhood. **g**, HomoDSP better matches the uPBM-derived C/G-favoring profile. Template attribution highlights residues that help disambiguate locally similar protein-DNA environments with different nucleotide preferences.

Taking position 10 as an example, DeepPBS predicted a weak T-over-C preference (Fig. 6c), whereas the uPBM-derived profile supported a C/G-favoring preference (Fig. 6g). We then retrieved training-set positions similar to DBP48 position 10 using a local protein-DNA interface descriptor and the DeepPBS hidden representation immediately before PWM readout. Both analyses showed T-biased neighborhoods, with the hidden-space retrieval showing particularly marked checkpoint-dependent T bias (Fig. 6d). These results suggest that DeepPBS maps the DBP48 position-10 environment to training examples in which similar local protein-DNA configurations are associated with T-favoring labels.

A representative hidden-space neighbor was `1hf0_A` position 9, which carries a strongly T-biased PWM label with *P* (*T*) = 0.960 and *P* (*C*) = 0.031 (Fig. 6e). DBP48 aligned to the `1hf0_A` training complex over 63 DBD residues, and DeepPBS attribution highlighted nearby basic and polar residues around DBP48 position 10 (Fig. 6f). Thus, the erroneous T-over-C prediction appears to be associated with local-interface similarity to T-favoring training examples.

These observations highlight an ambiguity of single-structure DBS prediction: similar local protein-DNA environments can be associated with different nucleotide preferences. DeepPBS, which lacks explicit homologous template information, appears to interpret the DBP48 position-10 environment as a T-favoring local pattern, despite the experimentally measured C/G preference.

HomoDSP alleviated this ambiguity by incorporating homologous template-interface features. At DBP48 position 10, HomoDSP produced a profile closer to the uPBM-derived C/G preference, whereas DeepPBS remained biased toward T (Fig. 6g). Template attribution showed that the correction was supported by residues from structurally matched templates, indicating that HomoDSP uses homologous interface evidence to distinguish locally similar but specificity-divergent protein-DNA environments. Together, this case supports the utility of template-enhanced geometric modeling for correcting nucleotide-level errors caused by ambiguous local contacts.

## Methods

### The predicted homologous complex database

We constructed a predicted homologous protein-DNA complex database by integrating BioLiP2 DBP annotations, homologous sequence retrieval, DNA fragment pairing, and Protenix-based complex structure prediction.

#### BioLiP2 DBP extraction and redundancy reduction

DBP entries were extracted from the BioLiP2 DNA subset [25]. For each record, we retained the PDB/chain ID, protein sequence, and DNA-binding-site annotation. The initial BioLiP2 DBP profile contained 24,456 chains. After removing exact sequence duplicates, these sequences were further clustered using DIAMOND `linclust` [26]. Clustering was performed at 70% sequence identity with 80% member coverage to obtain representatives for homologous DBD retrieval.

#### DBD inference and homologous sequence retrieval

For each 70% representative protein, DBD windows were inferred from the indices of DNA-binding residues. Each window spanned the contacting residue interval with 80-residue flanks and was capped at 500 amino acids; distant contacting regions separated by more than 250 residues were split into separate windows. Each window was searched against the combined UniRef50 [27] and BFD-Small [28] databases using DIAMOND. Hits were filtered by sequence identity, query coverage, and coverage of annotated contacting residues, and the top 100 bit-score-ranked hits per window were retained. Extracted homolog segments were capped at 500 amino acids.

#### Pairing homologous proteins with DNA fragments

For each DBD window, the corresponding DNA sequence was extracted from BioLiP2 DNA chain annotations or recovered from the matched `.mmCIF` file. DNA fragments between 6 and 80 nucleotides were used directly. For longer DNA chains, protein-contacting DNA regions were identified from the complex structure using a 6.0 Å atom-distance cutoff, extended by 10 nucleotides on both sides, and capped at 80 nucleotides. DNA records with unreliable cropping or missing sequence information were excluded from structure prediction. The resulting DBD-DNA fragment pairs were used as inputs for homologous complex structure prediction.

#### Homologous complex structure prediction

Each homologous DBD segment was paired with the DNA fragment associated with its originating DBD window. Protein MSAs were generated using `MMseqs2-GPU` [29] against the UniRef50/BFD-Small database. Each Protenix input contained one homologous DBD chain and one paired DNA chain. Complex structures were predicted using the Protenix default model (`protenix_base_default_v0.5.0`, https://github.com/bytedance/Protenix). Excluding failed cases, we obtained 143,724 predicted homologous protein-DNA complex structures (named HomoDB-FULL). 47,677 samples with ipTM > 0.4 and pTM > 0.5 were selected to form the high-quality template library (named HomoDB-HQ).

### Template-enhanced geometric neural networks

HomoDSP extends the geometric neural network architecture of DeepPBS by incorporating homologous protein-DNA interface information into the local DNA readout. The baseline model represents the query protein as an atom-level geometric graph and the DNA as an ordered set of base-pair positions with local DNA-shape descriptors. Protein-DNA interactions are encoded through geometric cross-message passing between protein atoms and DNA points, followed by a one-dimensional convolutional readout that predicts a four-channel PWM at each DNA position. In this setting, the query DNA sequence is not used as an input feature.

To provide homologous structural context, HomoDSP augments this query-only representation with a template-derived interface embedding. For each query complex, structurally similar protein-DNA interfaces are retrieved from HomoDB-FULL/HQ. For each query DNA position, HomoDSP identifies the corresponding position in each retrieved template and collects nearby template protein residues to form a local interface neighborhood. These residues are described by features capturing amino-acid identity, physicochemical properties, residue-DNA contact geometry, spatial distance, local orientation, and structural-alignment context.

A dedicated template-interface encoder summarizes the residue-level neighborhoods into position-specific template embeddings. Within each template, local residues around a DNA position are aggregated to represent the homologous interface environment. Multiple templates are then combined according to their structural matching scores, producing a single template-derived embedding for each query DNA position. This embedding is concatenated with the original protein-conditioned DNA representation and DNA-shape descriptors before the final convolutional PWM readout.

This design preserves the sequence-independent query setting of DeepPBS while adding residue-level homologous interface evidence. By conditioning each DNA position on structurally matched template residues, HomoDSP enables the network to use conserved protein-DNA contact patterns and local geometric compatibility to resolve nucleotide-specific preferences. For a detailed description of the feature representation and network structure, see Appendix A.

### Evaluation datasets

This study employed two datasets for evaluation.

- **The 5-fold CV benchmark dataset:** This dataset was collected and processed by DeepPBS from JASPAR2022 and contains 523 samples, each comprising a protein-chain-DNA complex and a corresponding PWM label (https://doi.org/10.6084/m9.figshare.25678053). We adopted the 5-fold CV split defined by DeepPBS and performed 5-fold cross-validation.
- **The 33 newly released JASPAR2026 samples**. We constructed an external test set from the JASPAR2026 CORE non-redundant motifs to assess generalization. We downloaded 2,633 CORE non-redundant PFMs and extracted UniProt accessions. Entries with multiple accessions were split into individual identifiers and mapped to experimentally resolved PDB structures. We then removed redundancy with the DeepPBS 5-fold CV benchmark. Candidate JASPAR2026 entries were first filtered by explicit overlap, including duplicated PDB entries, sample identifiers, and motif-structure assignments. To further prevent structural leakage, each remaining candidate was compared against all 523 benchmark structures using Foldseek [30], and candidates whose maximum TM-score against any benchmark sample *>* 0.9 were discarded. This explicit and structural de-redundancy yielded 39 candidates. Due to the structural constraints of the DeepPBS/X3DNA [31] preprocessing pipeline, including the requirement for a valid protein-DNA interface and a well-defined DNA helix, 33 of the 39 de-redundant JASPAR2026 candidates were successfully converted into DeepPBS-compatible feature files and corresponding HomoDSP template-augmented feature files. The final JASPAR2026 external test set contains 33 protein-DNA specificity samples from 21 unique PDB structures. Homologous template features were obtained by Foldseek retrieval and converted into HomoDSP top-16 template features. This external set was used only for evaluation and was not used for model selection or hyperparameter tuning.

### Implementation details

In this study, training and inference for the HomoDSP model were implemented using the PyTorch framework. A batch size of 1 was used. The Adam optimizer was employed with a learning rate of 0.001 and a weight decay of 0.0001. All experiments were conducted on a single NVIDIA RTX 4090 GPU. The model was trained for 50 epochs. The total number of trainable parameters is 17,152, of which 1,251 belong to the template encoder.

#### Training loss

Let **P** ∈ ℝ^*L×*4^ denote the ground-truth PWM and 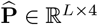 denote the predicted PWM. A binary mask 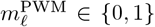 indicates whether DNA position *ℓ* has a valid PWM label. The model is trained with a masked PWM reconstruction loss:

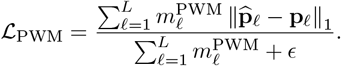

An additional information-content consistency term is applied to discourage overly diffuse or overly sharp PWM predictions. The overall objective is

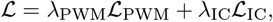

where *λ*_PWM_ and *λ*_IC_ control the relative contributions of PWM reconstruction and information-content consistency.

### Gradient-based interpretability analysis

We used integrated gradients implemented in Captum [32] to quantify which input features contributed to a selected PWM prediction. For a PWM position *j*, we defined the attribution target as a logit contrast between two nucleotides,

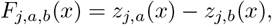

where *z*_*j,a*_(*x*) and *z*_*j,b*_(*x*) denote the model logits for nucleotide *a* and a competing nucleotide *b* at position *j*, respectively. This contrastive target attributes the model evidence supporting nucleotide *a* over nucleotide *b* at the selected PWM column, thereby avoiding aggregation of attribution signals across unrelated PWM positions.

Integrated gradients were computed with respect to the selected input tensor *x*, using a zero tensor *x*^*′*^ of the same shape as the baseline. For feature dimension *i*, the attribution was defined as

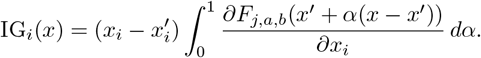

In practice, the integral was approximated using a finite Riemann sum,

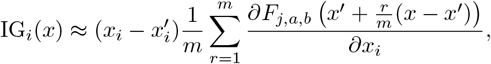

where *m* denotes the number of interpolation steps. Unless otherwise stated, absolute integrated-gradient values were used for ranking and aggregation, whereas signed attributions were retained when the direction of contribution was analyzed.

For the no-template baseline, attributions were computed with respect to the protein atom graph and DNA-shape inputs. The protein attribution tensor assigns a contribution score to each atom-level node and each atom feature. Atom-level scores were summarized as

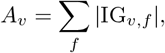

where *v* denotes an atom node and *f* denotes an atom-level feature. Residue-level scores were then obtained by summing atom-level scores over all atoms assigned to the same residue,

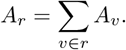

This aggregation produced both atom-level and residue-level explanations for the selected PWM column. The same principle was used to summarize attributions over DNA positions or DNA-shape feature groups.

For HomoDSP, template-specific attribution was computed with respect to the template-interface representation while keeping the query-complex inputs fixed. The attributed tensor contains template-interface features indexed by template rank *k*, DNA-context position *l*, template residue node *m*, and feature channel *f*. The feature channels include amino-acid identity and physicochemical properties, atom–DNA context features, residue-DNA distance and contact features, distance radial-basis encodings, local geometric descriptors, and alignment-derived features. Invalid template residue nodes were excluded using the template-node mask before aggregation.

The attribution score for a template residue node was calculated as

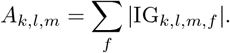

Template-level, DNA-position-level, and feature-block-level scores were obtained by summing over the corresponding axes:

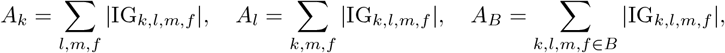

where *B* denotes a predefined feature block, such as atom–DNA context, distance/contact features, local geometry, distance radial-basis features, or alignment-derived features. When a specific template residue was selected, its feature-level attribution vector,

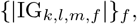

was reported directly to identify which geometric or biochemical descriptors contributed most strongly to the nucleotide contrast. For ensemble predictions, attribution was computed independently for each checkpoint and averaged after the same aggregation procedure.

### uPBM data processing

We used uPBM data from the study by Glasscock *et al*. [13], which reports 7-mer E-scores and Z-scores for DBP1, DBP3, DBP5, DBP6, DBP9, DBP35, and DBP48. For each design, all DNA 7-mers with valid E-scores were extracted from the replicate-specific sheets. Each 7-mer was paired with its reverse complement, and the lexicographically smaller sequence was used as the canonical 7-mer identifier. Replicate measurements and reverse-complement-equivalent entries were then averaged, yielding one mean uPBM E-score for each canonical 7-mer. These mean E-scores were used as 7-mer-level experimental binding labels for model evaluation.

For a predicted PWM of length *L*, we scanned all possible 7-bp windows. For each window and each canonical 7-mer *s*, the model score was defined as the larger log-probability between the forward sequence and its reverse complement:

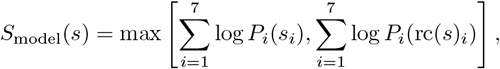

where *P*_*i*_(*b*) denotes the predicted probability of nucleotide *b* at position *i* within the scanned window. Model scores were compared with mean uPBM E-scores across all canonical 7-mers using Spearman correlation, PCC, and AUROC. Since the predicted PWM and the uPBM 7-mer measurements may differ in positional register, the 7-bp window with the highest Spearman correlation was selected as the best-matching window for each protein and model.

The uPBM data were not used for model training, model selection, or full-length PWM supervision of these AI-designed proteins. For visualization only, we derived an empirical uPBM logo by selecting the top-scoring uPBM 7-mers and accumulating base frequencies at each of the seven positions, weighted by shifted E-scores. The resulting matrix was normalized column-wise and used only as a qualitative representation of the experimentally enriched 7-mer preference.

## Conclusion

In this study, we present HomoDSP, a template-enhanced framework for DBS prediction from protein-DNA complex structures. By constructing a large-scale database of predicted homologous protein-DNA complexes, HomoDSP expands specificity modeling beyond individual natural complexes and enables the use of diverse homologous interface patterns for DBS prediction. The benchmark, external validation, AF3-structure, and uPBM analyses consistently support the value of this homologous recognition landscape. Template attribution analyses further indicate that HomoDSP uses DNA-contacting residues and structure-aware interface features, supporting both the effectiveness and interpretability of the framework. Its performance on AF3-predicted complexes and AI-designed DNA-binding proteins suggests a scalable route from sequence to predicted structure to DBS prediction.

Despite these advances, HomoDSP has several limitations. First, the framework depends on an external homologous template library, and its performance is influenced by the availability and quality of predicted protein-DNA complexes. Although confidence filtering and template attribution analyses support the utility of the current database, low-confidence or specificity-divergent templates may still introduce noise for certain families or samples. Second, HomoDSP still requires a protein-DNA complex structure as input. Therefore, its application is currently most suitable for proteins with experimentally resolved complexes or reliable AF3-like predicted structures; improving the full sequence-to-structure-to-specificity workflow remains an important direction. Third, HomoDSP was not specifically designed to model single-mutation effects. Because template retrieval is primarily driven by overall structural and interface similarity, the current framework may have limited sensitivity to subtle specificity changes caused by individual amino-acid substitutions. Systematic evaluation on mutation-resolved protein-DNA binding datasets will be required to determine and improve its ability to predict mutation-induced specificity shifts.

## Data availability

All raw data in this study were obtained from publicly available databases. The source code and 33 JASPAR2026 samples are freely available at https://github.com/wwzll123/HomoDSP. The benchmark dataset for five-fold cross-validation can be downloaded from https://doi.org/10.6084/m9.figshare.25678053. The predicted homologous complex database is deposited at Zenodo https://doi.org/10.5281/zenodo.20675636.

## Acknowledgements

This work was supported by NSFC Grants 625B2068; NSFC-FDCT Grant 62361166662; National Key R&D Program of China 2023YFC3503400 and 2022YFC3400400; The Innovative Research Group Project of Hunan Province 2024JJ1002; Hunan Science and Technology Innovation Plan 2025ZYJ003; Key R&D Program of Hunan Province 2023GK2004, 2023SK2059, and 2023SK2060; Top 10 Technical Key Project in Hunan Province 2023GK1010; Key Technologies R&D Program of Guangdong Province (2023B1111030004 to FFH); and Postgraduate Research Innovation Project of Hunan Province CX20250637. We would like to thank the Fund of the National Supercomputing Center in Changsha (http://nscc.hnu.edu.cn/), Peng Cheng Lab, Key Laboratory of High-Performance Distributed Ledger Technology and Digital Finance (Ministry of Education), and Hunan Research Center of the Basic Discipline for Cell Signaling.

## Appendix A Network architecture details

### Notation and input representation

For a query protein-DNA complex, the protein is represented as an atom-level graph

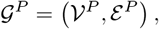

where *V*^*P*^ denotes the set of protein atoms and *ε*^*P*^ denotes protein graph edges defined by covalent connectivity and local spatial neighborhoods. Each protein atom *i* ∈ *V*^*P*^ is associated with a 3D coordinate 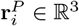, an atom feature vector 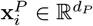, and a local orientation vector 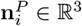. The atom feature vector contains atom-type and physicochemical descriptors, including charge, atomic radius, solvent accessibility, Atchley-factor descriptors, and local coordination features.

The DNA is represented as an ordered sequence of *L* base-pair positions,

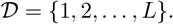

For each DNA position *ℓ* ∈ *D*, a set of coarse-grained DNA points is used to describe local DNA chemical groups:

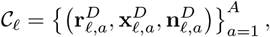

where *A* is the number of coarse-grained DNA points per base-pair position, 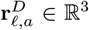 is the coordinate of the *a*-th DNA point, 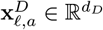 is its feature vector, and 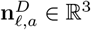 is its local orientation vector.

Each DNA position is also associated with a DNA-shape descriptor

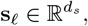

where *d*_*s*_ = 14 in this work. The shape descriptor contains six intra-base-pair parameters,

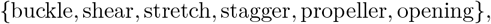

six inter-base-pair parameters,

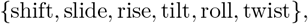

and two groove-width descriptors corresponding to major- and minor-groove widths. The nucleotide sequence of the query DNA is not used as an input feature.

### Query protein-DNA geometric encoder

Protein atom features are first encoded by a protein graph encoder:

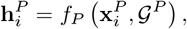

where *f*_*P*_ denotes the protein graph neural network and 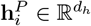 is the encoded representation of atom *i*.

To model local protein-DNA interactions, a bipartite graph is constructed between protein atoms and DNA coarse-grained points. A protein atom *i* and a DNA point (*ℓ, a*) are connected if their distance is below a cutoff *r*_*c*_:

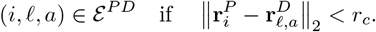

For each protein–DNA edge, the relative displacement vector is

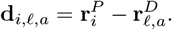

A geometric edge descriptor is then defined as

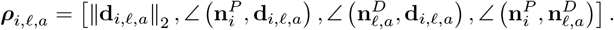

The protein-conditioned representation of each DNA point is computed by geometric message passing:

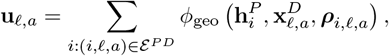

where *ϕ*_geo_ is a learnable multilayer perceptron. Messages are computed from local 3D geometry and can be separated according to DNA chemical groups, such as phosphate, sugar, and base-associated points.

The DNA-point representations at position *ℓ* are then reduced to a base-pair-level representation:

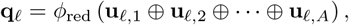

where ⊕ denotes feature concatenation and *ϕ*_red_ is a learnable projection network. The vector **q**_*ℓ*_ represents the query protein-conditioned DNA environment at position *ℓ*.

### Template-interface representation

For each query complex, HomoDSP retrieves up to *K* homologous protein-DNA template complexes from the predicted template library. Let *T*_*k*_ denote the *k*-th retrieved template and let *s*_*k*_ denote its structural matching score.

For each query DNA position *ℓ*, the corresponding DNA position in template *T*_*k*_ is determined from the template DNA mapping. Around this mapped template position, nearby protein residues are collected to form a local template-interface neighborhood:

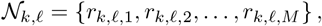

where *M* is the maximum number of retained residues. A binary mask

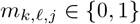

indicates whether the *j*-th residue is present in the local neighborhood.

Each template residue *r*_*k,ℓ,j*_ is represented by a residue-level feature vector:

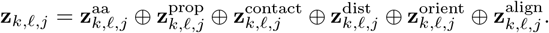

Here, **z**^aa^ encodes amino-acid identity, and **z**^prop^ encodes physicochemical residue classes, including hydrophobic, polar, positively charged, negatively charged, aromatic, aliphatic, small, sulfur-containing, glycine, and proline categories. The contact feature **z**^contact^ describes the closest protein atom, whether it belongs to the backbone or side chain, and the contacted DNA chemical group. The distance feature **z**^dist^ describes residue–DNA spatial proximity. The orientation feature **z**^orient^ describes residue geometry relative to the local DNA coordinate frame. The alignment feature **z**^align^ encodes structural alignment context between the query and template interfaces.

Distance features include minimum residue–DNA distances, group-specific distances to phosphate, sugar and base atoms, local contact counts, and radial basis function expansions. For a distance value *d*, the *b*-th radial basis feature is defined as

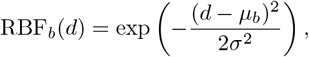

where *µ*_*b*_ is the center of the *b*-th radial basis and *σ* controls its width.

To describe local orientation, a DNA-centered coordinate frame is constructed at each mapped template DNA position:

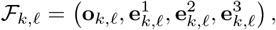

where **o**_*k,ℓ*_ is the local origin and 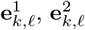, and 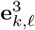 are orthonormal axes derived from the DNA geometry. A residue coordinate **r**_*k,ℓ,j*_ can be expressed in this local frame as

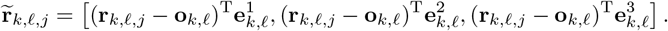

Similar local-frame projections are used for residue-to-DNA direction vectors and side-chain orientation vectors.

### Template-interface encoder

The template-interface encoder is the main component that transfers homologous interface evidence into the PWM predictor. Its input is the residue neighborhood *N*_*k,ℓ*_ around the DNA position in each matched template, together with the structural matching score *s*_*k*_ and the validity mask *m*_*k,ℓ,j*_. The encoder is designed to be residue-aware, position-specific and template-score weighted, so that the model can use homologous protein–DNA contacts without directly using the nucleotide sequence of the query DNA.

Each residue-level feature vector is first projected into a common hidden space:

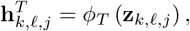

where 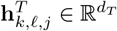.

For a fixed template *T*_*k*_ and query DNA position *ℓ*, the valid residue embeddings are then summarized into a local interface descriptor. In practice, the implementation combines masked statistical pooling with a learned residue-importance score, but these operations are used only to obtain a compact representation of the local protein environment and are not treated as separate model assumptions. The resulting per-template interface embedding is written as

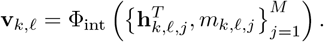

This descriptor preserves the amino-acid composition, residue–DNA contact pattern, distance profile, and local-frame orientation of the homologous interface around position *ℓ*.

Multiple templates may provide evidence for the same query position. Their contributions are weighted by the structural matching scores:

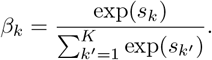

A position-specific validity indicator removes templates that do not provide any residue neighbor for the mapped DNA position:

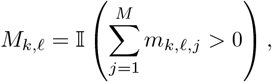

where *M*_*k,ℓ*_ = 1 if template *T*_*k*_ provides at least one valid residue for position *ℓ*, and *M*_*k,ℓ*_ = 0 otherwise. The final template-derived embedding at DNA position *ℓ* is therefore

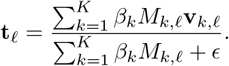

Thus, **t**_*ℓ*_ summarizes homologous residue-level interface information from all retrieved templates while down-weighting weak structural matches and ignoring unmapped positions. If no valid template evidence is available at position *ℓ*, **t**_*ℓ*_ is set to a zero vector and the model falls back to the query-only geometric branch.

### Template-enhanced PWM readout

For each DNA position *ℓ*, HomoDSP combines three sources of information: the query protein-conditioned DNA representation **q**_*ℓ*_, the DNA-shape descriptor **s**_*ℓ*_, and the template-derived interface embedding **t**_*ℓ*_. The fused representation is

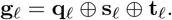

A one-dimensional convolutional readout then models short-range dependencies along the DNA axis and maps **g**_1:*L*_ to the predicted PWM 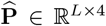. The readout is intentionally kept lightweight so that the template channel contributes local interface evidence rather than replacing the query protein–DNA geometric encoder.

To account for DNA strand symmetry, the model is applied to both the original DNA orientation and the reverse-complement orientation. The two orientation-specific predictions are used jointly during training and evaluation.

## Appendix B Evaluation Metrics

We used the following metrics to evaluate the agreement between predicted and reference nucleotide preferences. Unless otherwise stated, metrics were computed over all evaluated PWM positions and nucleotide channels.

### Mean absolute error

Mean absolute error (MAE) measures the average absolute deviation between the predicted value and the reference value. For *N* evaluated entries, with prediction ŷ_*i*_ and reference value *y*_*i*_, MAE is defined as

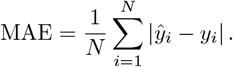

A lower MAE indicates better agreement with the reference values.

### Root mean squared error

Root mean squared error (RMSE) measures the square-root of the average squared prediction error,

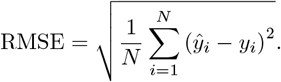

Compared with MAE, RMSE gives larger penalties to large individual errors. A lower RMSE indicates better agreement.

### Pearson correlation coefficient

The Pearson correlation coefficient (PCC) measures the linear correlation between predicted and reference values,

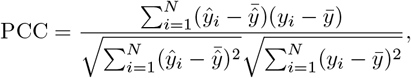

where 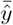 and 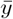 are the means of the predicted and reference values, respectively. PCC ranges from −1 to 1, with larger values indicating stronger positive linear agreement.

### Spearman rank correlation

Spearman correlation measures the monotonic agreement between predicted and reference values after converting both variables to ranks. Let *R*(*ŷ*_*i*_) and *R*(*y*_*i*_) denote the ranks of *ŷ*_*i*_ and *y*_*i*_. Spearman correlation is computed as the Pearson correlation between these ranks:

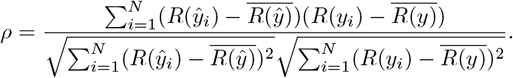

Larger *ρ* values indicate better agreement in the relative ordering of sequence preferences.

### Area under the receiver operating characteristic curve

The area under the receiver operating characteristic curve (AUROC) evaluates how well a continuous prediction score separates positive and negative labels. For a threshold *t*, the true positive rate and false positive rate are

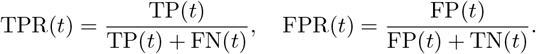

AUROC is the area under the curve obtained by plotting TPR(*t*) against FPR(*t*) over all possible thresholds:

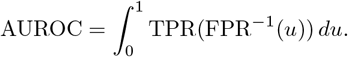

Equivalently, AUROC is the probability that a randomly selected positive example receives a higher prediction score than a randomly selected negative example. A value of 0.5 corresponds to random ranking, and a value of 1.0 indicates perfect separation.

### Top-base accuracy

Top-base accuracy evaluates whether the nucleotide with the highest predicted probability matches the nucleotide with the highest reference probability at each PWM position. For position *j*, let

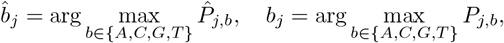

where 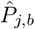 and *P*_*j,b*_ are the predicted and reference probabilities for nucleotide *b* at position *j*. For *L* evaluated PWM positions, top-base accuracy is

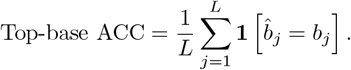

This metric measures positional agreement in the most preferred nucleotide, independent of the full probability distribution at each column.

**Figure A1.**
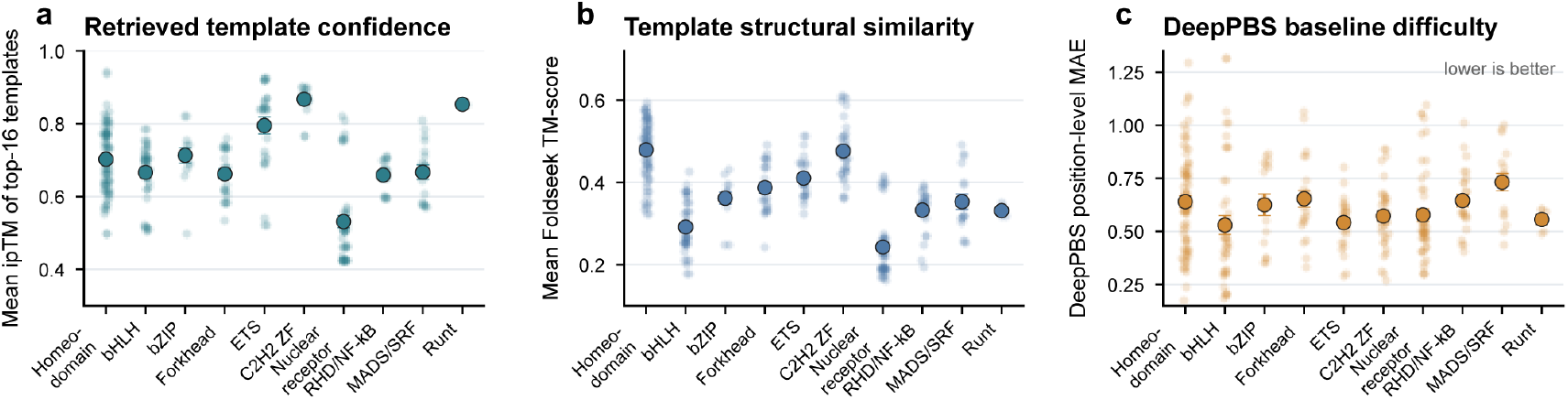
Template quality grouped by family and baseline error.

## References

(1) Lambert, S. A.; Jolma, A.; Campitelli, L. F.; Das, P. K.; Yin, Y.; Albu, M.; Chen, X.; Taipale, J.; Hughes, T. R.; Weirauch, M. T. The human transcription factors. Cell 2018, 172, 650–665.

(2) Trujillo-Ochoa, J. L.; Kazemian, M.; Afzali, B. The role of transcription factors in shaping regulatory T cell identity. Nature Reviews Immunology 2023, 23, 842–856.

(3) He, Z.; He, J.; Xie, K. KLF4 transcription factor in tumorigenesis. Cell Death Discovery 2023, 9, 118.

(4) Zeng, W.; Dou, Y.; Pan, L.; Xu, L.; Peng, S. Improving prediction performance of general protein language model by domain-adaptive pretraining on DNA-binding protein. Nature Communications 2024, 15, 7838.

(5) Lambert, S. A.; Yang, A.; Sasse, A.; Cowley, G.; Albu, M.; Caddick, M.; Morris, Q.; Weirauch, M.; Hughes, T. Similarity regression predicts evolution of transcription factor sequence specificity. Nature Genetics 2019, 51, 981–989.

(6) Wetzel, J. L.; Zhang, K.; Singh, M. Learning probabilistic protein-DNA recognition codes from DNA-binding specificities using structural mappings. Genome Research 2022, 32, 1776–1786.

(7) Liu, S.; Gomez-Alcala, P.; Leemans, C.; Glassford, W. J.; Melo, L. A. N.; Lu, X.-J.; Mann, R. S.; Bussemaker, H. J. Predicting the DNA binding specificity of transcription factor mutants using family-level biophysically interpretable machine learning. Nucleic Acids Research 2025, 53, gkaf831.

(8) Ovek Baydar, D.; Rauluseviciute, I.; Aronsen, D. R.; Blanc-Mathieu, R.; Bonthuis, I.; De Beukelaer, H.; Ferenc, K.; Jegou, A.; Kumar, V.; Lemma, R. B., et al. JASPAR 2026: expansion of transcription factor binding profiles and integration of deep learning models. Nucleic acids research 2026, 54, D184–D193.

(9) Butcher, J. et al. De novo Design of All-atom Biomolecular Interactions with RFdiffusion3. bioRxiv 2025, DOI: 10.1101/2025.09.18.676967.

(10) Geraseva, E.; Bokov, G.; Polovnikov, V.; Golovin, A. Leveraging Auxiliary Potentials in RFDiffusion for the Design of NA-Binding Proteins. Journal of Chemical Information and Modeling 2026, 66, 4463–4471.

(11) Sehgal, E.; Politanska, Y.; Mitra, R.; Kim, P. T.; Gonzalez Rodriguez, N.; Warrier, T.; Kubaney, A.; Morishita, A.; Quijano, R.; Butcher, J., et al. Generative design of sequence specific DNA binding proteins. bioRxiv 2026, 2026–04.

(12) Ichikawa, D. M. et al. A universal deep-learning model for zinc finger design enables transcription factor reprogramming. Nature Biotechnology 2023, 41, 1117–1129.

(13) Glasscock, C. J. et al. Computational design of sequence-specific DNA-binding proteins. Nature Structural & Molecular Biology 2025, 32, 2252–2261.

(14) Mitra, R.; Li, J.; Sagendorf, J. M.; Jiang, Y.; Cohen, A. S.; Chiu, T.-P.; Glasscock, C. J.; Rohs, R. Geometric deep learning of protein-DNA binding specificity. Nature Methods 2024, 21, 1674–1683.

(15) Kubaney, A.; Favor, A.; McHugh, L.; Mitra, R.; Pecoraro, R.; Dauparas, J.; Glasscock, C.; Baker, D. RNA sequence design and protein–DNA specificity prediction with NA-MPNN. bioRxiv 2025, 2025–10.

(16) Abramson, J.; Adler, J.; Dunger, J.; Evans, R.; Green, T.; Pritzel, A.; Ronneberger, O.; Willmore, L.; Ballard, A. J.; Bambrick, J., et al. Accurate structure prediction of biomolecular interactions with AlphaFold 3. Nature 2024, 630, 493–500.

(17) Weirauch, M. et al. Determination and inference of eukaryotic transcription factor sequence specificity. Cell 2014, 158 6, 1431–1443.

(18) Zenker, S.; Wulf, D.; Meierhenrich, A.; Viehöver, P.; Becker, S.; Eisenhut, M.; Stracke, R.; Weisshaar, B.; Bräutigam, A. Many transcription factor families have evolutionarily conserved binding motifs in plants. Plant Physiology 2024, 198, DOI: 10.1093/plphys/kiaf205.

(19) Shen, N.; Zhao, J.; Schipper, J.; Zhang, Y.; Bepler, T.; Leehr, D.; Bradley, J.; Horton, J.; Lapp, H.; Gordân, R. Divergence in DNA Specificity among Paralogous Transcription Factors Contributes to Their Differential In Vivo Binding. Cell systems 2018, 6 4, 470–483.e8.

(20) Team, P.; Zhang, Y.; Gong, C.; Zhang, H.; Ma, W.; Liu, Z.; Chen, X.; Guan, J.; Wang, L.; Yang, Y., et al. Protenix-v1: Toward high-accuracy open-source biomolecular structure prediction. bioRxiv 2026, 2026–02.

(21) Laforet, M.; McMurrough, T. A.; Vu, M.; Brown, C. M.; Zhang, K.; Junop, M.; Gloor, G.; Edgell, D. Modifying a covarying protein-DNA interaction changes substrate preference of a site-specific endonuclease. Nucleic Acids Research 2019, 47, 10830–10841.

(22) Watson, L. C.; Kuchenbecker, K.; Schiller, B.; Gross, J.; Pufall, M.; Yamamoto, K. R. The glucocorticoid receptor dimer interface allosterically transmits sequence-specific DNA signals. Nature structural & molecular biology 2013, 20, 876–883.

(23) Hastings, R.; Aditham, A.; DelRosso, N.; Suzuki, P.; Fordyce, P. Mutations to transcription factor MAX allosterically increase DNA selectivity by altering folding and binding pathways. Nature Communications 2025, 16, DOI: 10.1038/s41467-024-55672-2.

(24) Zhang, C.; Shine, M.; Pyle, A. M.; Zhang, Y. US-align: universal structure alignments of proteins, nucleic acids, and macromolecular complexes. Nature methods 2022, 19, 1109–1115.

(25) Zhang, C.; Zhang, X.; Freddolino, L.; Zhang, Y. BioLiP2: an updated structure database for biologically relevant ligand–protein interactions. Nucleic acids research 2024, 52, D404–D412.

(26) Buchfink, B. J.; Barbé, É.; Ashkenazy, H.; Reuter, K.; Kennedy, J. A.; Drost, H.-G. Clustering the protein universe of life using DIAMOND DeepClust. Nature Methods 2026, 724–727.

(27) Suzek, B. E.; Wang, Y.; Huang, H.; McGarvey, P. B.; Wu, C. H.; the UniProt Consortium UniRef clusters: a comprehensive and scalable alternative for improving sequence similarity searches. Bioinformatics 2015, 31, 926–932.

(28) Jumper, J. et al. Highly accurate protein structure prediction with AlphaFold. Nature 2021, 596, 583–589.

(29) Kallenborn, F.; Chacon, A.; Hundt, C.; Sirelkhatim, H.; Didi, K.; Cha, S.; Dallago, C.; Mirdita, M.; Schmidt, B.; Steinegger, M. GPU-accelerated homology search with MMseqs2. Nature Methods 2025, 22, 2024–2027.

(30) Van Kempen, M.; Kim, S. S.; Tumescheit, C.; Mirdita, M.; Lee, J.; Gilchrist, C. L.; Söding, J.; Steineg-ger, M. Fast and accurate protein structure search with Foldseek. Nature biotechnology 2024, 42, 243–246.

(31) Lu, X.-J.; Bussemaker, H. J.; Olson, W. K. DSSR: an integrated software tool for dissecting the spatial structure of RNA. Nucleic Acids Research 2015, 43, e142.

(32) Kokhlikyan, N.; Miglani, V.; Martin, M.; Wang, E.; Alsallakh, B.; Reynolds, J.; Melnikov, A.; Kliushk-ina, N.; Araya, C.; Yan, S.; Reblitz-Richardson, O. Captum: A unified and generic model interpretability library for PyTorch. arXiv 2020, 2009.07896.

